# Robustness measures for cell type classification in single-cell RNA-seq datasets of kidney organoids

**DOI:** 10.64898/2026.09.20.752971

**Authors:** Harel Reinuss, Jacob Goldberger, Tomer Kalisky

## Abstract

**Background:** Induced pluripotent stem cell (iPSC)-derived kidney organoids hold great promise for disease modeling, drug screening, and regenerative medicine. However, since these organoids are synthetic constructs, it is essential to evaluate how faithfully their constituent cell populations recapitulate the corresponding cell types in the native kidney. Single-cell transcriptomic label-transfer tools, which assign cell-type labels by integrating cells with well-annotated reference data and classifying them, are increasingly used for this purpose. Yet the reliability of the resulting labels — particularly for immature or partially differentiated cells that might have no clear in vivo counterpart — has not been systematically addressed.

**Methodology:** To quantify the reliability of cell-type labels assigned by label-transfer algorithms, we introduce two complementary robustness measures for each cell type — the *classification stability score*, which assesses label stability under simulated Poisson noise, and the *prediction specificity score*, which assesses how well the predicted cell-type label probabilities match the assigned labels. We apply this methodology to scRNA-seq datasets from the human fetal kidney and four iPSC-derived kidney organoid protocols, using a mouse fetal kidney dataset as a reference.

**Results:** We find that label-transfer robustness scores are generally high in most human fetal kidney cell types, except for cells of immature epithelial structures. In the organoid-derived cell subpopulations, robustness scores are high for the cap mesenchyme but vary for epithelial lineages according to each protocol’s specific differentiation strategy. These observations are consistent across both classification stability and prediction specificity scores, which are generally well correlated with each other. We further validate our approach through a simulated cell-removal experiment, confirming that the proposed measures correctly identify unreliable classifications.

**Significance:** These robustness measures provide a generalizable framework for assessing the reliability of cell-type label transfer in single-cell transcriptomic datasets, and can be extended to other biomedical settings that rely on automated single-cell classification.

## INTRODUCTION

In recent years, numerous protocols have been developed for culturing organoids from induced pluripotent stem cells (iPSCs) for various tissues including the brain [1], colon [2], mammary gland [3], and kidney [4]. These organoids can be used for disease modeling, personalized drug testing, and regenerative medicine. For the fetal kidney, which is the focus of this study, various protocols have been developed [5] with particular emphasis on differentiation into specific cell types such as the collecting duct [6], podocytes [7], or the proximal tubule [8].

Because iPSC-derived organoids are synthetic constructs, it is essential to evaluate the degree to which they mimic the desired tissue [9]. For example, a fetal kidney organoid should ideally contain cells representing the various cell compartments of the fetal kidney, that is, the nephrogenic-zone stroma (the un-induced metanephric mesenchyme), the cap mesenchyme (containing nephron progenitor cells, NPCs), early epithelial structures (comma and S-shaped bodies), and differentiated epithelial structures such as the podocytes, the proximal tubule, the loop of Henle, the distal tubule, and the collecting duct [10].

To date, the capacity of an organoid to differentiate into desired cell types is typically evaluated using tissue staining and immunofluorescence for specific gene markers, functional tests, and single-cell RNA-seq to assess the expression of gene sets characteristic of each cell type. More recently, classification tools such as DevKidCC [11] have been developed to perform automated cell-type labeling using human fetal kidney scRNA-seq data as a reference (Figure 1A).

**Figure 1:**
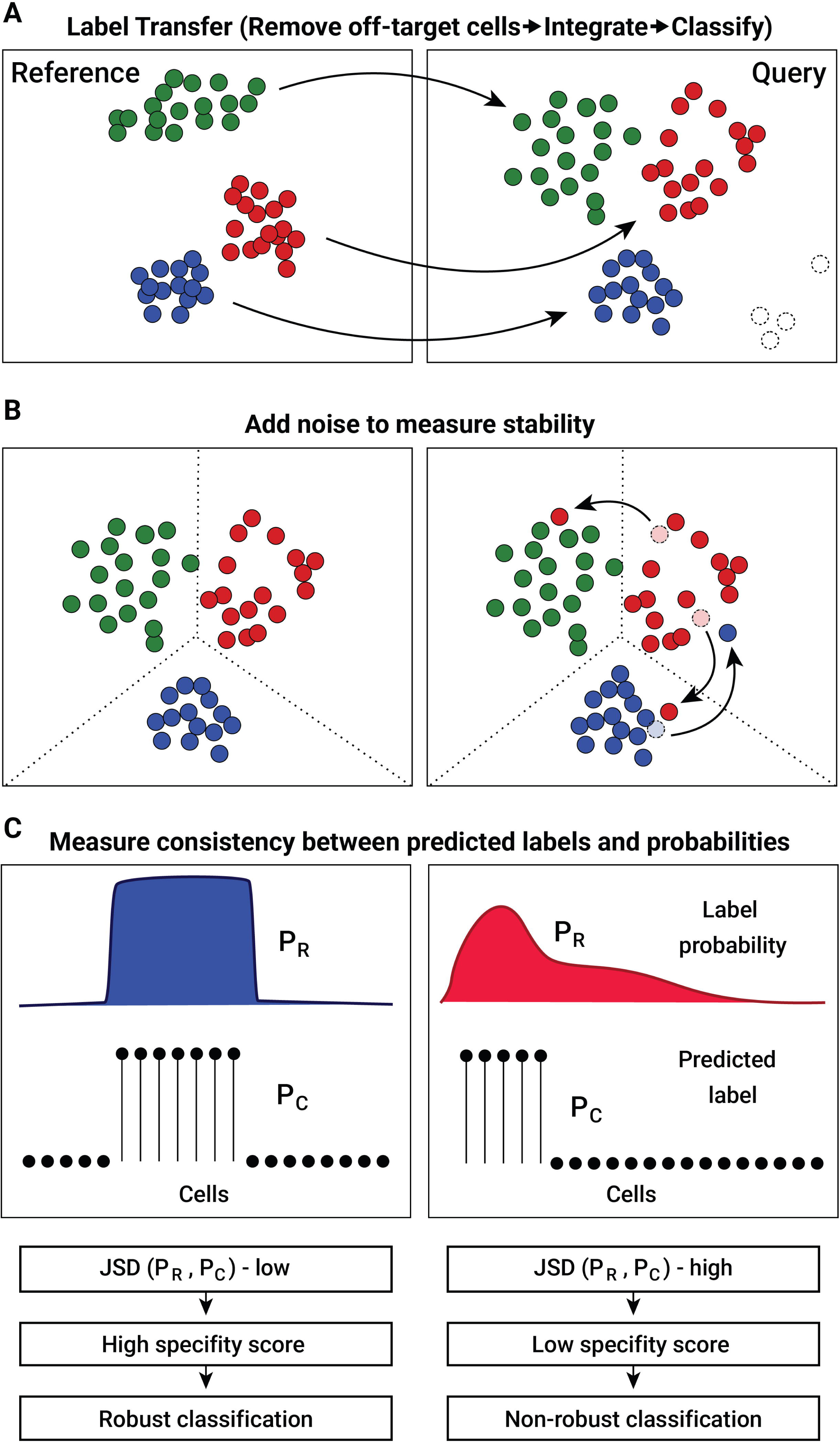
A methodology for estimating label-transfer robustness in single-cell transcriptomic datasets using classification stability and prediction specificity scores. (A) After removing non-renal lineage (off-target) cells from the query dataset, we perform label transfer, which consists of integrating the query and reference datasets and then classifying the query cells based on the reference labels. Cell labels are represented by colors. Off-target cells are shown as dashed circles. (B) The classification stability score is estimated by introducing Poisson noise into the query dataset and repeating the label transfer process. The stability score of a cell subpopulation quantifies the combined fraction of cells that, following the introduction of noise, either lost their label (moved to another subpopulation) or gained it incorrectly (moved in from another subpopulation) relative to the cell subpopulation’s original size. (C) The prediction specificity score quantifies the discrepancy between the label-transfer algorithm’s final decision regarding a cell’s label (*P_C_*) and the predicted probability for that label (*P_R_*). This reflects the degree to which the label-transfer algorithm is confident in its decision.

Cell type labeling in kidney organoids presents several challenges. First, organoids frequently contain non-renal lineage (off-target) cell subpopulations, such as melanocytes, neurons, and muscle cells [6], that must be distinguished from kidney-lineage cells. These off-target cells are relatively easy to identify, since their transcriptomic profiles differ markedly from kidney cell types. For example, DevKidCC exploits this by using PAX2 expression as a kidney-specific lineage filter [11]. A more fundamental challenge is evaluating how faithfully kidney-like subpopulations recapitulate their in vivo counterparts. Immature or incompletely differentiated cells may display mixed or partial phenotypes that have no clear equivalents in the tissue-derived reference data, making confident label assignment difficult. Finally, even in the reference data, cell type annotations are based on specific marker genes rather than “ground truth”, thereby introducing uncertainty that propagates into any classification built upon them.

In this study, we evaluate the reliability of cell-type labels assigned by single-cell label-transfer algorithms. These algorithms typically perform data integration [12,13] followed by classification (Figure 1A). We argue that an assigned label is reliable if it is robust, that is, stable under small perturbations introduced by technical or biochemical noise. Conversely, cells with mixed or partial phenotypes are likely to have transcriptomic profiles located near the boundary between cell types, resulting in low-confidence label assignments that are unstable and may flip to a different cell type under small amounts of noise.

Therefore, to assess the reliability of assigned cell-type labels, we propose two complementary measures of label-transfer robustness. For each cell type we calculate the *classification stability score*, which measures the degree to which assigned cell-type labels are conserved after the addition of Poisson noise (Figure 1B), and the *prediction specificity score*, which measures how well the predicted cell-type label probabilities match the assigned labels (Figure 1C). We demonstrate our methodology in scRNA-seq datasets from the human fetal kidney and four iPSC-derived kidney organoid protocols, using a mouse fetal kidney dataset as a reference.

## RESULTS

### A methodology for estimating label-transfer robustness using classification stability and prediction specificity scores

To demonstrate our methodology we used a reference scRNA-seq dataset from a mouse fetal kidney [14], which was manually labeled according to known marker genes (Figure 2A), and used Seurat’s ‘MapQuery’ algorithm [12] to perform label transfer onto a human fetal kidney dataset from the kidney cell atlas [15] (Figure 2B). The reference dataset contains the following cell subpopulations: the uninduced mesenchyme or nephrogenic zone stroma (UM), the cap mesenchyme (CM), the proliferating cap mesenchyme (CM_DIV), early epithelial structures such as C/S-shaped bodies (PROX_1), podocytes (PODO), fetal proximal tubule (PROX_2), loop of Henle (LOH), distal tubule and collecting duct (DIST_CD), macrophages (MACROPHAG), and endothelium (ENDO), all of which could also be well discerned in the query dataset.

**Figure 2:**
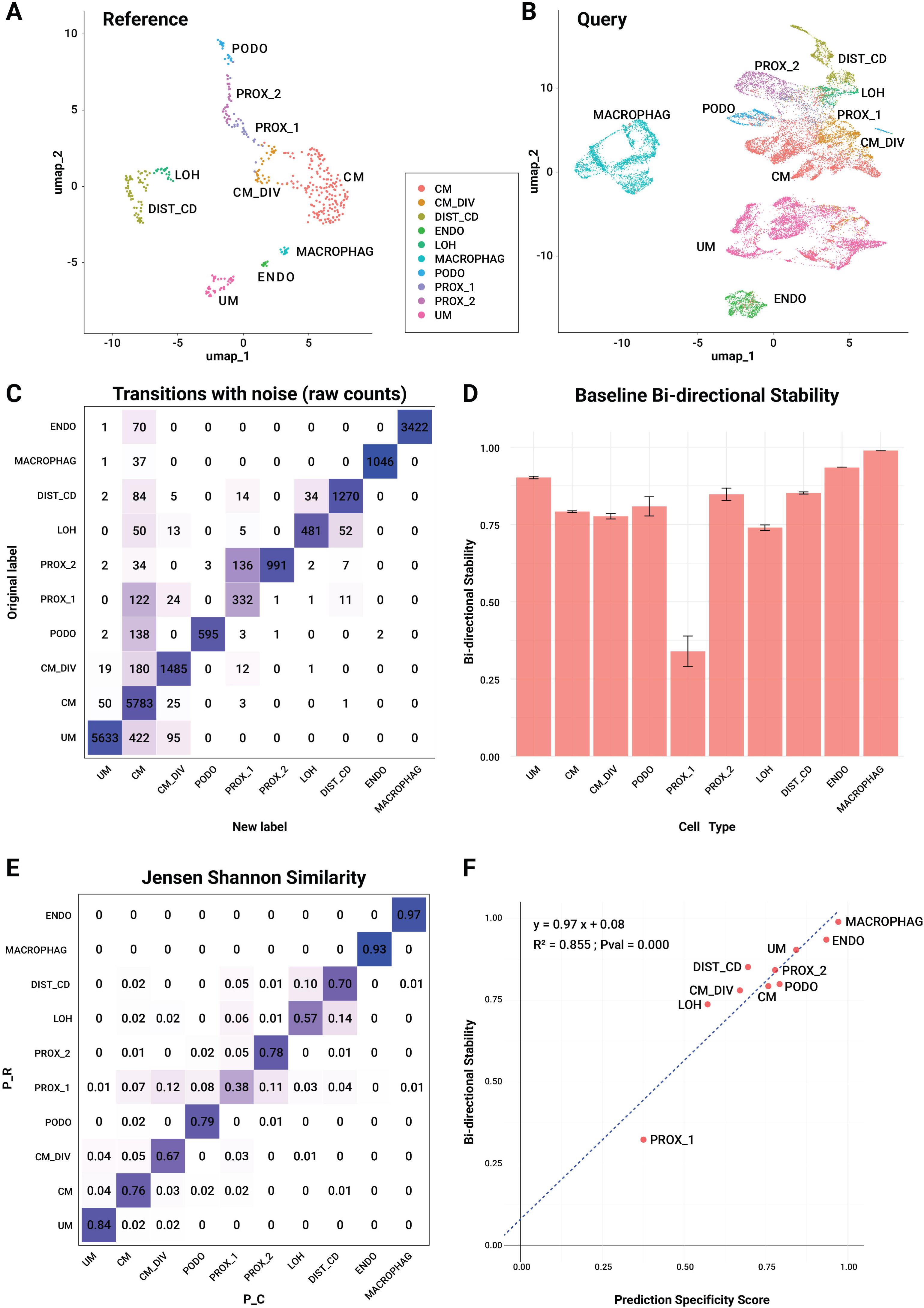
Cell-label transfer from the mouse fetal kidney to the human fetal kidney shows high classification stability and prediction specificity scores across cell subpopulations, except for cells of immature epithelial structures. (A) A UMAP plot of the single-cell RNA-seq reference dataset from the mouse fetal kidney. Cell labels were taken from the original publication, in which cells were manually annotated based on known marker genes. (B) A UMAP plot of the single-cell RNA-seq query dataset from the human fetal kidney (kidney cell atlas). Cells are colored according to their predicted labels. (C) A transition matrix showing the number of cells that switched labels after Poisson noise was applied. Diagonal entries are colored according to the stability score 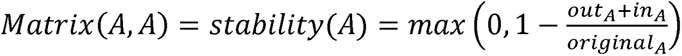. Off-diagonal entries are colored according to their effect on both the origin and destination subpopulations: 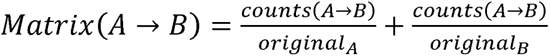. Classification stability scores of all cell subpopulations, averaged over N=10 noise realizations. (E) A matrix of Jensen-Shannon similarity scores, defined as 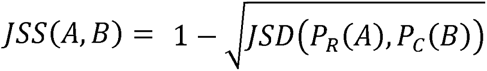. Color intensity reflects the similarity score for each cell-type pair. (F) A scatter plot of classification stability versus prediction specificity scores for all cell subpopulations. The two measures show a strong correlation. Cell type abbreviations: UM - the uninduced mesenchyme or nephrogenic zone stroma, CM - the cap mesenchyme, CM_DIV - the proliferating cap mesenchyme, PROX_1 - early epithelial structures such as C/S-shaped bodies, PODO - podocytes, PROX_2 - fetal proximal tubule, LOH - loop of Henle, DIST_CD - distal tubule and collecting duct, MACROPHAG - macrophages, ENDO - endothelium.

To quantify the **classification stability** score for each cell type, we introduced Poisson noise into the query dataset by resampling each gene expression count from a Poisson distribution with mean equal to the original count, and then repeated the label transfer procedure. We define the “bi-directional stability” of a predicted cell subpopulation as one minus the fraction of cells that changed their labels as a result of the noise:

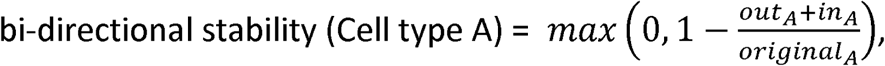

where: *Out_A_* is the number of cells that changed their label from cell type A to another cell type, *in_A_* is the number of cells that changed their label from another cell type to cell type A, and *original_A_* is the total number of cells labeled as cell type A before the addition of noise (Figures 1B and 2C-D). This score penalizes cell subpopulations containing many cells whose cell-type label changes because of Poisson noise. Hereafter, we will refer to the bi-directional stability as “stability”.

As a complementary measure, we calculated the **prediction specificity score** for each predicted cell type (Figure 1C and 2E), which we defined as the Jensen-Shannon similarity [16–18]:

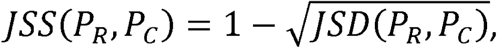

Where: 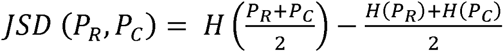 is the Jensen-Shannon divergence between the label probabilities 0 ≤ *P_R_* ≤ 1 and the final predicted labels *P_C_* = 0 *or* 1, and *H*(*P*) = − Σ*_i_ P_i_ log*(*P_i_*) is the entropy. This score measures how well the final cell-type decision matches the probability calculated by the label-transfer algorithm, that is, the degree to which the label-transfer algorithm is confident in its decisions.

We found that these two scores correlated with each other (*R*^2^ = 0.855, Figure 2F). Both are relatively high for all human fetal kidney cell subpopulations profiled in the kidney cell atlas, except for the cell subpopulation containing early epithelial structures (PROX_1), whose low scores reflect its content of incompletely differentiated - and therefore mixed and poorly defined - phenotypes.

### In the iPSC-derived organoids, label-transfer robustness scores are high for the cap mesenchyme but vary for epithelial lineages

We next calculated classification stability and prediction specificity scores for publicly available scRNA-seq datasets from organoids generated by the Takasato et al. protocol [4] (dataset from [6]) (Figure 3A-C), the Uchimura et al. protocol [6] (Figure 3D-F), the Harder et al. protocol [7] (Figure S1), and the Vanslambrouck et al. protocol [8] (D13 and D13+14, Figure S2). In all organoids we observed a high correlation between classification stability and prediction specificity scores (R² = 0.67–0.88, Figures 2F, 3C, 3F, S1C, S2C, S2F). Likewise, all organoid datasets had high stability scores for either the cap mesenchyme or the proliferating cap mesenchyme (maximum of CM and CM_DIV stability = 0.65–0.90, Table S1).

**Figure 3:**
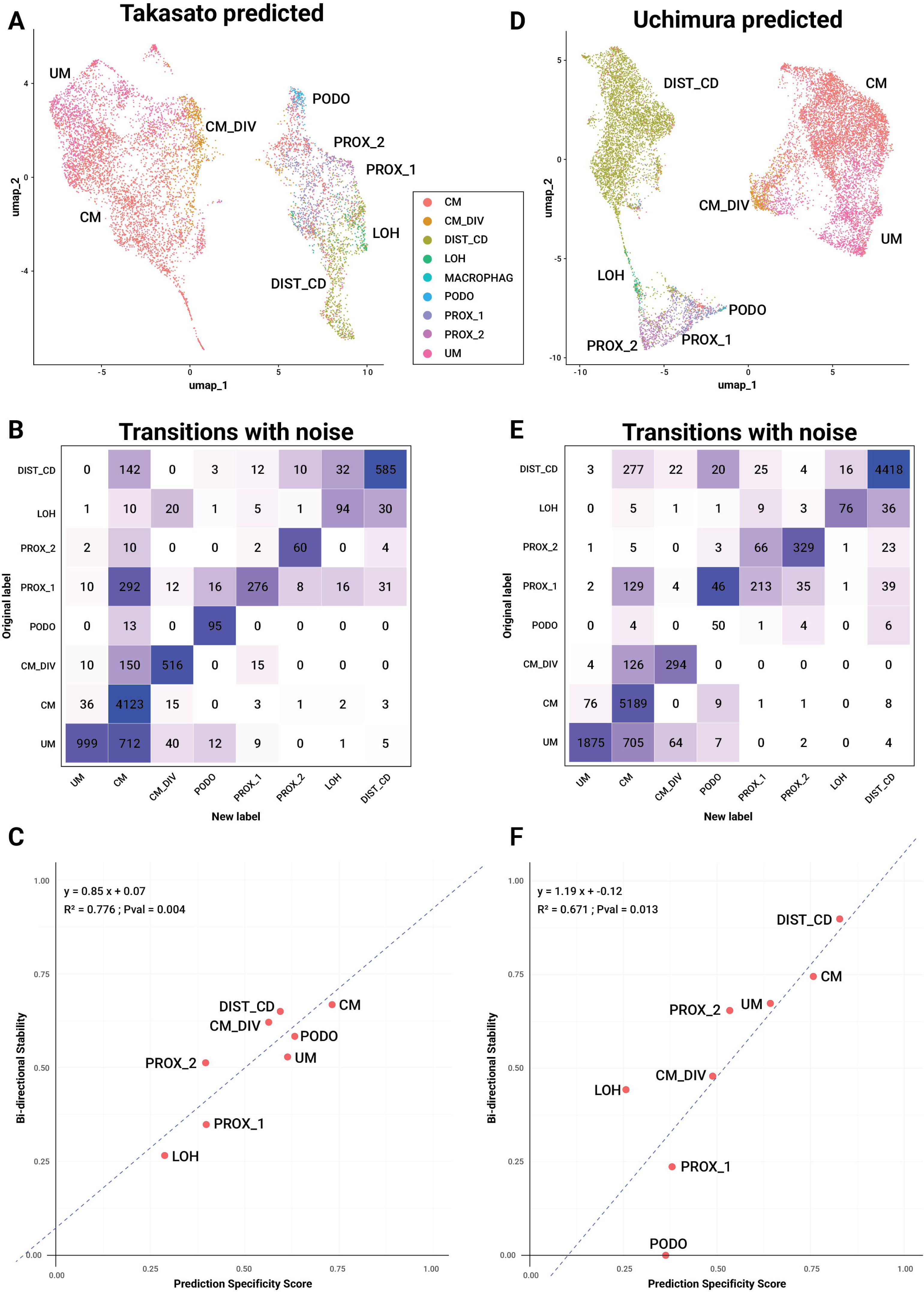
Cell-label transfer from the mouse fetal kidney to the Takasato et al. and Uchimura et al. iPSC-derived kidney organoids shows a robustly classified cap mesenchyme and varying robustness for epithelial lineages. (A,D) UMAP plots of the single-cell RNA-seq query datasets from the Takasato et al. and Uchimura et al. iPSC-derived kidney organoids. Cells are colored according to their predicted labels. (B,E) Transition matrices showing the number of cells that switched labels after Poisson noise was applied. Matrix entries are colored as in Figure 2. (C,F) Scatter plots of classification stability versus prediction specificity scores for all cell subpopulations. The two measures show a strong correlation in both organoids. Both protocols have high robustness scores for the cap mesenchyme lineage (CM, stability = 0.67 for Takasato et al. and 0.75 for Uchimura et al.), but varying scores for the epithelial lineages. For example, the Takasato et al. organoid has a higher score for podocytes (PODO, stability = 0.58), whereas the Uchimura et al. organoid has a higher score for distal tubule and collecting duct lineages (DIST_CD, stability = 0.90).

Regarding the epithelial cell subpopulations, the stability scores varied between the different organoid protocols and cell subpopulations (Table S1). Specifically, the **Takasato** et al. organoid had the highest score for the early epithelial structures (PROX_1, stability = 0.34), which was also comparable to the human fetal kidney - consistent with this protocol’s design for recapitulating both nephron progenitor cells and early nephron differentiation. Note that the early epithelial cell subpopulation (PROX_1) consistently scored low for all organoids as well as for the human fetal kidney (stability = 0.12–0.34). The **Uchimura** et al. organoid exhibited high scores for the distal tubule and collecting duct cell subpopulation (DIST_CD, stability = 0.90), in line with this protocol’s design to favor collecting duct lineages. Similarly, the **Harder** et al. organoid had high scores for podocytes (PODO, stability = 0.80), consistent with the high content of cells classified as “early glomerular epithelia” or “maturing podocytes” in the original study. The **Vanslambrouck** et al. dataset included two time points: the D13 monolayer stage in which most epithelial lineages are not yet present, and the D13+14 stage, which had a high score for the fetal proximal tubule (PROX_2, stability = 0.91), consistent with the enhanced proximal tubular compartment reported in the original study. Note that the D13+14 stage also scored high for the nephrogenic zone stroma cell subpopulation (UM, stability = 0.89), consistent with the large stromal component reported in the original study.

### Simulated removal of cell subpopulations from a query dataset results in decreased classification stability scores

To validate our methodology, we reasoned that if a cell subpopulation is removed from the query dataset (but not from the reference dataset), the label-transfer algorithm will nonetheless re-integrate all remaining query cells with the reference and, as a result, might still assign the absent subpopulation’s label to other cells from neighboring subpopulations with similar transcriptomic profiles — producing unreliable cell labels. If our methodology is correct, these cells should have low classification stability scores.

To test this, we took the human fetal kidney scRNA-seq dataset from the kidney cell atlas, whose cells had been previously labeled using the mouse fetal kidney as a reference, and artificially removed specific cell subpopulations before repeating the label transfer procedure. When the loop of Henle (LOH), early epithelial structures (PROX_1), and distal tubule and collecting duct (DIST_CD) cell subpopulations were removed (Figure 4B), the label-transfer algorithm still assigned these labels to some cells in the query dataset, but with low classification stability scores. This finding indicates that the stability score correctly flags unreliable label assignments.

**Figure 4:**
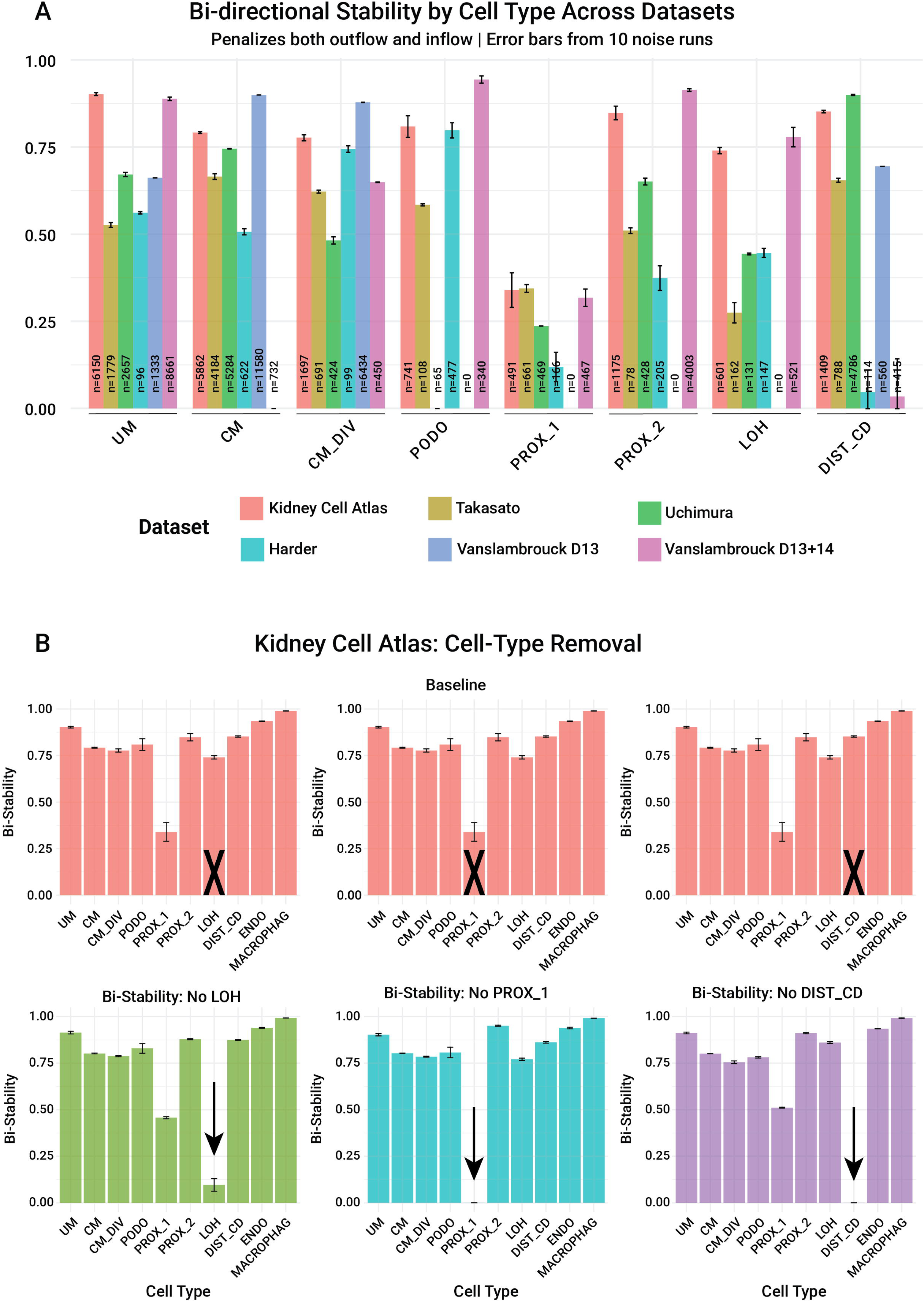
Simulated removal of cell subpopulations from a query dataset results in decreased classification stability scores. (A) Classification stability scores for all cell subpopulations from the human fetal kidney and iPSC-derived kidney organoid datasets. All organoid datasets have high stability scores for either the cap mesenchyme (CM) or the proliferating cap mesenchyme (CM_DIV), whereas the epithelial cell subpopulations (PODO, PROX_1, PROX_2, LOH, and DIST_CD) have stability scores that vary between different organoid protocols and cell subpopulations. (B) After simulated removal of specific cell subpopulations from the human fetal kidney dataset (kidney cell atlas), the label-transfer algorithm still assigns these labels to some of the remaining cells in the query, but with low stability scores that indicate unreliable labeling. All scores were averaged over N=10 noise realizations.

## DISCUSSION

In this study, we propose robustness measures that quantify the reliability of cell-type labels assigned by label-transfer algorithms to single-cell transcriptomic profiles.

Automated classification of cells based on single-cell transcriptomes and other high-dimensional features is likely to become widely used in biomedical applications - for example, assessing the immune exhaustion state of T-cells or macrophages within the tumor microenvironment. In such settings, robustness measures such as those proposed in this study could help evaluate the reliability of the assigned cell-type or cell-state labels and flag unreliable ones.

We chose the mouse fetal kidney [14] as our reference despite two potential limitations: cross-species differences that could introduce bias, and a relatively small dataset size (∼600 cells). We nonetheless considered it a suitable reference for several reasons. First, it was sequenced using the Smart-seq2 protocol, which provides deep, full-length transcript coverage and low dropout rates, yielding more reliable expression profiles for individual cells. Second, the major cell subpopulations were well represented, clearly separated, and reliably annotated. Third, at the resolution of the 10 major cell types used in our classification, the transferred labels yielded cell subpopulations in the human fetal kidney whose marker gene expression resembled that of their mouse reference counterparts (Figure S3). Most importantly, classification of the Uchimura et al. organoid dataset using the human fetal kidney dataset (from the kidney cell atlas) and its cell-type annotations as the reference yielded similar results to those obtained when using the mouse fetal kidney as reference (Figure 3F for mouse reference; Figure S4F for human reference), with high robustness scores for collecting duct lineages in both cases (“DIST_CD” and “CNT/PC proximal UB”, respectively) and a strong correlation between classification stability and prediction specificity scores (R² = 0.67 and 0.81, respectively). Note that some cell subpopulations of early epithelial structures (e.g., proximal, medial, and distal S-shaped body) have high scores when using the human fetal kidney dataset as reference, likely due to the finer resolution of its cell-type annotations.

For the classification stability score, we used a bi-directional measure that penalizes the stability of cell type A based on cells that changed their labels following the addition of Poisson noise in either direction - both from the original label A to another label and from other labels to label A. This is similar to the “total flip rate” measure used to evaluate the similarity of the results produced by two alternative classifiers (that is, the original and noisy).

We also tested two additional classification stability scores. The first measures the fraction of cells that retained their original label after noise was added, and penalizes only cells that changed their label from the original cell type A to another cell type:

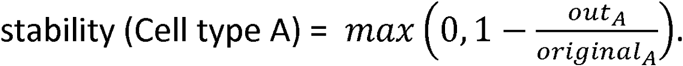

The second is an F1-based stability score, defined as the harmonic mean of precision 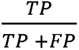 and recall 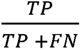.

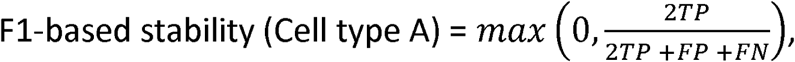

where: *TP* = the number of cells that retained their original labels, *FP*= *in_A_* = the number of cells that changed their label from other cell type to cell type A, and *FN*= *out_A_* = the number of cells that changed their label from cell type A to another cell type. Neither score was as strongly correlated with the prediction specificity score as the bi-directional stability score.

A major challenge in estimating the reliability of classification algorithms for labeling single-cell transcriptomic profiles is the absence of “ground truth”. This challenge is particularly pronounced for organoid cells, which are not sampled from native tissue, but are instead artificially differentiated in culture using defined growth factors and timing protocols. To address this limitation, we treated the initial (noise-free) classification as a substitute for ground truth and measured how consistently the classifier’s results were reproduced when noise was introduced, rather than evaluating them against an independent standard.

An additional challenge in estimating the reliability of single-cell transcriptomic profile classification algorithms is the need to integrate datasets collected from different laboratories, organisms (e.g., human vs. mouse), biological systems (e.g., native tissue vs. organoid), and experimental protocols (e.g., organoid generation protocols, single-cell library preparation methods). Here, we used Seurat’s MapQuery() function to integrate and classify the data. Notably, our robustness scoring methodology is not specific to this integration and classification tool, and could, in principle, be applied to evaluate the reliability of other cell labeling and classification tools, such as DevKidCC [11].

We note that a more general noise model would use the negative binomial distribution, which has been shown to more accurately capture biological (overdispersed) noise in scRNA-seq data [19,20]. However, this would require estimating an additional dispersion parameter, thereby increasing model complexity. We leave this extension for future work.

## METHODS

### Data preprocessing and quality control

#### The mouse fetal kidney reference dataset (Smart-seq2)

The gene expression counts matrix was downloaded from GEO accession code GSE146988 [14]. Non-viable and low-quality cells – specifically, those labeled as low-quality in the original publication or demonstrating zero counts for housekeeping genes (Actb/Gapdh) – were filtered out. Genes that were expressed in fewer than 10 cells were excluded.

#### The human fetal kidney dataset from the kidney cell atlas (10x)

The gene expression counts matrix (“Fetal_full_v3.h5ad”) was downloaded from the kidney cell atlas website (https://www.kidneycellatlas.org/). Non-renal cell subpopulations were excluded based on the metadata annotations in the original publication [15]. Specifically, we removed: B cells, T cells (CD4+ and CD8+), natural killer cells, dendritic cells, neurons, erythroid cells, megakaryocytes, mast cells, neutrophils, and fibroblasts.

#### The Takasato et al. organoid dataset (10x)

The gene expression counts matrix was downloaded from GEO accession code GSM3763146 [6]. After initial UMAP projection and clustering, non-renal lineage (off-target) cells were identified based on known marker genes: MLANA for melanocytes, MAP2 and SOX2 for neurons, and COL1A1 for putative non-renal stromal cells. Marker gene expression was visualized using Seurat’s FeaturePlot function, and off-target cell subpopulations were removed using Seurat’s CellSelector function.

#### The Uchimura et al. organoid dataset (10x)

The gene expression counts matrix was downloaded from GEO accession code GSM3763147 [6]. After initial UMAP projection and clustering, non-renal lineage (off-target) cells were identified based on known marker genes: MLANA for melanocytes, SOX2 for neurons, and PITX2 for muscle cells. Marker gene expression was visualized using Seurat’s FeaturePlot function, and off-target cell subpopulations were removed using Seurat’s CellSelector function.

#### The Harder et al. organoid dataset (Drop-seq)

The gene expression counts matrix was downloaded from GEO accession code GSM3204322 [7]. Gene expression matrices from singly picked Day 20 organoids were filtered according to the quality control thresholds used in the original publication: cells were retained if they contained between 500 and 4,000 detected genes (nFeature_RNA) and a mitochondrial read percentage (percent.mt) below 25%.

#### The Vanslambrouck et al. organoid datasets (10x)

The gene expression counts for Day 13 and Day 13+14 were downloaded from GEO accession codes GSM5600482 and GSM5600483 [8], respectively, and imported using ReadMtx. Cells were retained if they contained between 500 and 9,000 detected genes (nFeature_RNA) and a mitochondrial read percentage (percent.mt) below 15%.

### Single-cell RNA-seq data processing and dimensionality reduction

Single-cell RNA-seq datasets were analyzed using the Seurat R package (v5) [21]. For each dataset, standard preprocessing was applied: raw count matrices were log-normalized using NormalizeData (scale factor 10,000), and the top 2,000 highly variable features were identified with FindVariableFeatures (selection.method = “vst”). Data were scaled with ScaleData, followed by Principal Component Analysis (RunPCA) for linear dimensionality reduction. Clustering was performed using FindNeighbors and FindClusters. UMAP embedding was performed using RunUMAP.

### Single-cell label transfer and mapping

Cell-type annotations were transferred from the reference dataset to the query dataset using Seurat’s anchor-based integration framework, with default parameters throughout. First, FindTransferAnchors was used to project the query dataset onto the reference’s principal component (PCA) space and identify anchor correspondences (mutual nearest neighbors) between reference and query cells. These anchors were then used by MapQuery, which is a wrapper around three sequential steps: (1) TransferData - transfers cell-type labels to the query cells, (2) IntegrateEmbeddings - uses the anchors to modify the query’s PCA projection to achieve an integrated embedding with the reference, and (3) ProjectUMAP - projects the query cells into the pre-computed reference UMAP coordinates, based on this integrated embedding.

### Noise perturbation and stability analysis

To evaluate the stability of transferred label predictions, synthetic noise was introduced to the query count matrices by resampling each gene expression count from a Poisson distribution with mean equal to the original count. This process was repeated for N=10 independent iterations (or N=5 for the analysis shown in Figure S4). Cell subpopulations within the query that contained fewer than 3 cells in the initial classification (before noise was introduced) were excluded from the stability analysis to prevent distortion due to small sample sizes.

## ACKNOWLEDGEMENTS

We wish to thank Ben Humphreys, Yehuda Neumark, and all the members of our lab for useful comments and suggestions.

## DECLARATION OF INTEREST STATEMENT

The authors have declared that no competing interests exist.

## AUTHOR CONTRIBUTIONS

Study initiation and conception – H.R. and T.K.; Data analysis - H.R. and T.K.; Other intellectual contribution – J.G.; Manuscript writing – H.R. and T.K.

## FUNDING

H.R. and T.K. were supported by the Israel Science Foundation (ICORE no. 1902/12 and Grants no. 1634/13, 2017/13, and 1814/20), the Israel Ministry of Health (Grant no. 3-10146), the EU-FP7 (Marie Curie International Reintegration Grant no. 618592), the Data Science Institute at Bar-Ilan University (seed grant), the ICRF (Grant no. 19-101-PG), the Israel Ministry of Science (Grant no. 3-16220), the Israel Ministry of Justice (Estates Committee / Va’adat HaIzavonot), and the Israel Cancer Association (Grant no. 20240114). The funders had no role in study design, data collection and analysis, decision to publish, or preparation of the manuscript.

## DECLARATION OF GENERATIVE AI AND AI-ASSISTED TECHNOLOGIES IN THE WRITING PROCESS

During the preparation of this work, the authors used Gemini (Google) and Claude (Anthropic) to discuss ideas, write and adapt code, and improve the readability and language of the text. After using these tools, the authors reviewed and edited all content and take full responsibility for the content of the publication.

## DATA AVAILABILITY STATEMENT

All datasets used in this study are publicly available: Mouse fetal kidney (Smart-seq2) - GEO accession code GSE146988; Kidney cell atlas, “fetal full” (10x) - kidney cell atlas website (https://www.kidneycellatlas.org/); Takasato et al. iPSC-derived kidney organoid (10x) - GEO accession code GSM3763146; Uchimura et al. iPSC-derived kidney organoid (10x) - GEO accession code GSM3763147; Harder et al. iPSC-derived kidney organoid (Drop-seq) - GEO accession code GSM3204322; and Vanslambrouck et al. iPSC-derived kidney organoid (10x) - GEO accession codes GSM5600482 and GSM5600483.

## CODE AVAILABILITY STATEMENT

The code used for analysis is available on GitHub at:

https://github.com/HaReL1/organoid-classifier-stability.

## KEY POINTS

▯ We propose two complementary robustness scores — classification stability and prediction specificity — for evaluating the reliability of cell-type labels assigned by label-transfer algorithms in single-cell transcriptomic datasets.
▯ The two measures were found to be well-correlated across multiple datasets tested.
▯ Cell-label transfer from the mouse fetal kidney to the human fetal kidney shows high robustness scores across most cell types, except for cells of immature epithelial structures.
▯ In the iPSC-derived kidney organoid datasets, label-transfer robustness scores are high for the cap mesenchyme but vary for epithelial lineages.
▯ These measures can help assess how well iPSC-derived organoids recapitulate specific cell types and may extend to other biomedical applications that rely on automated single-cell classification, such as evaluating exhaustion status of immune cells in a tumor microenvironment.

## APPENDICES

**Supplementary information:** Supplementary figures (Figures S1-S4).

**Supplementary Table S1:** Classification stability scores for all cell subpopulations in the human fetal kidney and iPSC-derived kidney organoid single-cell RNA-seq datasets, following cell-label transfer from the mouse fetal kidney dataset (provided as a separate spreadsheet file).

## Notes

### Competing Interest Statement

The authors have declared no competing interest.

