## Supplementary information: Supplementary figures (Figures S1-S4). for "Robustness measures for cell type classification in single-cell RNA-seq datasets of kidney organoids"

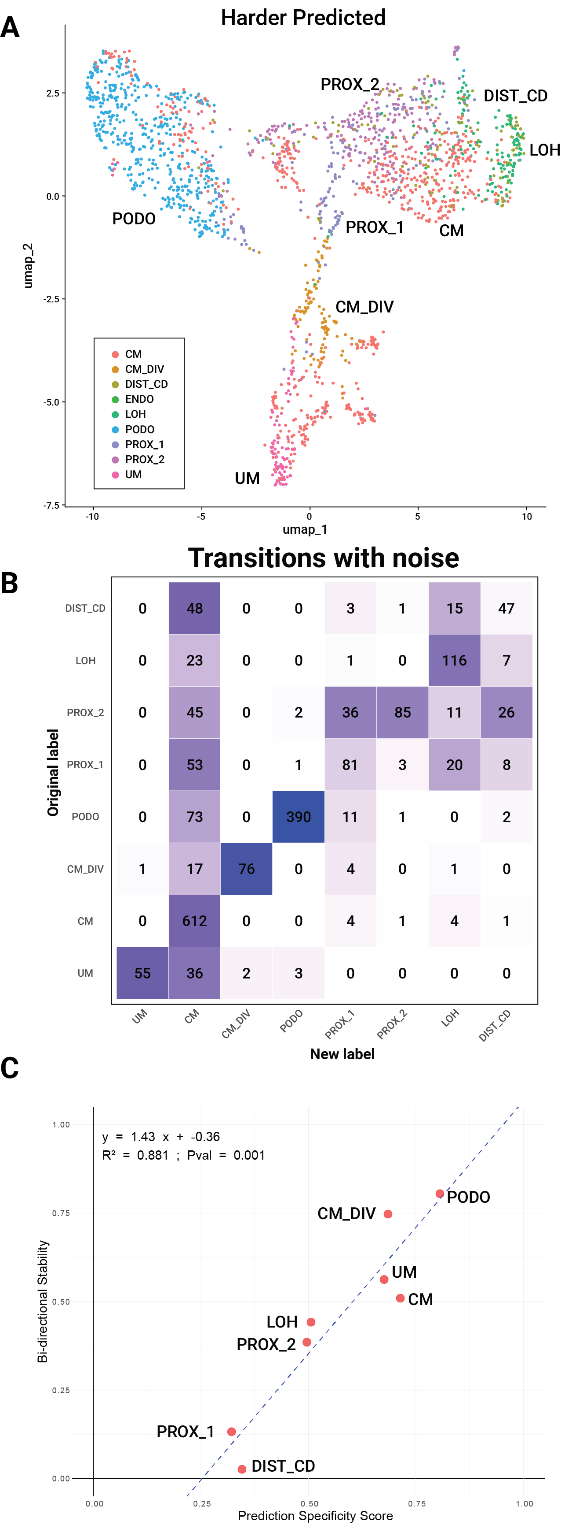

### Figure S1: Cell-label transfer from the mouse fetal kidney to the Harder et al. iPSC-derived kidney organoid shows a robustly classified cap mesenchyme and varying robustness for epithelial lineages.

(A) A UMAP plot of the single-cell RNA-seq query dataset from the Harder et al. iPSC-derived kidney organoid [1]. Cells are colored according to their predicted labels. (B) A transition matrix showing the number of cells that switched labels after Poisson noise was applied. Matrix entries are colored as in Figure 2. (C) A scatter plot of classification stability versus prediction specificity scores for all cell subpopulations. The two measures show a strong correlation. This protocol has high robustness scores for the cap mesenchyme (CM_DIV, stability = 0.74) and podocyte (PODO, stability = 0.80) lineages.

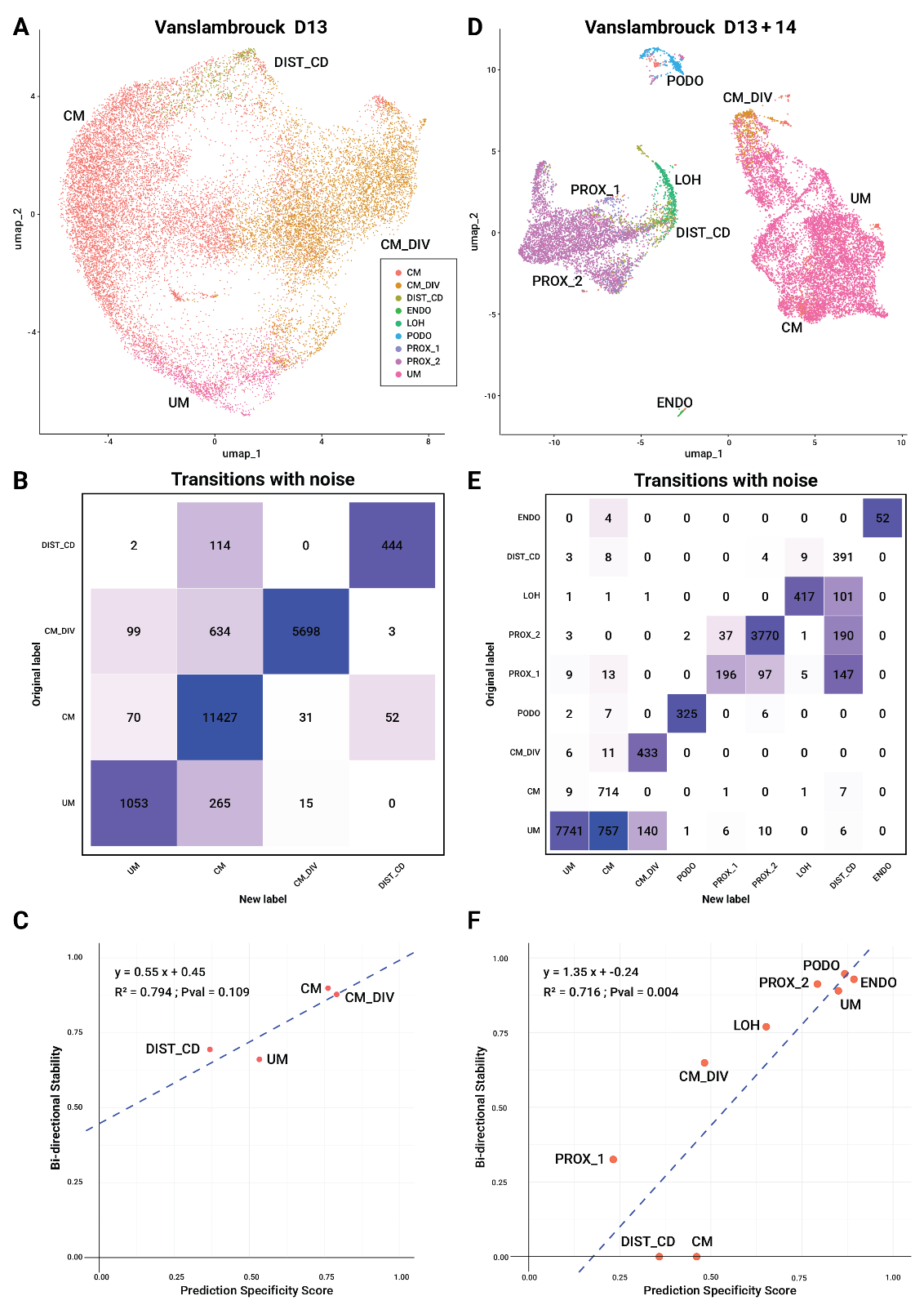

### Figure S2: Cell-label transfer from the mouse fetal kidney to the Vanslambrouck et al. iPSC-derived kidney organoids shows a robustly classified cap mesenchyme and varying robustness for epithelial lineages.

(A,D) UMAP plots of the single-cell RNA-seq query datasets from the Vanslambrouck et al. iPSC-derived kidney organoids, at D13 and D13+14 [2]. Cells are colored according to their predicted labels. (B,E) Transition matrices showing the number of cells that switched labels after Poisson noise was applied. Matrix entries are colored as in Figure 2. (C,F) Scatter plots of classification stability versus prediction specificity scores for all cell subpopulations. The two measures show a strong correlation in the D13+14 organoid. The D13 organoid has high robustness scores for the cap mesenchyme lineage (CM, stability = 0.90; CM_DIV, stability = 0.88). The D13+14 organoid has high robustness scores for the proliferating cap mesenchyme (CM_DIV, stability = 0.65), as well as for the nephrogenic zone stroma (UM, stability = 0.89), podocyte (PODO, stability = 0.94), and fetal proximal tubule (PROX_2, stability = 0.91) lineages.

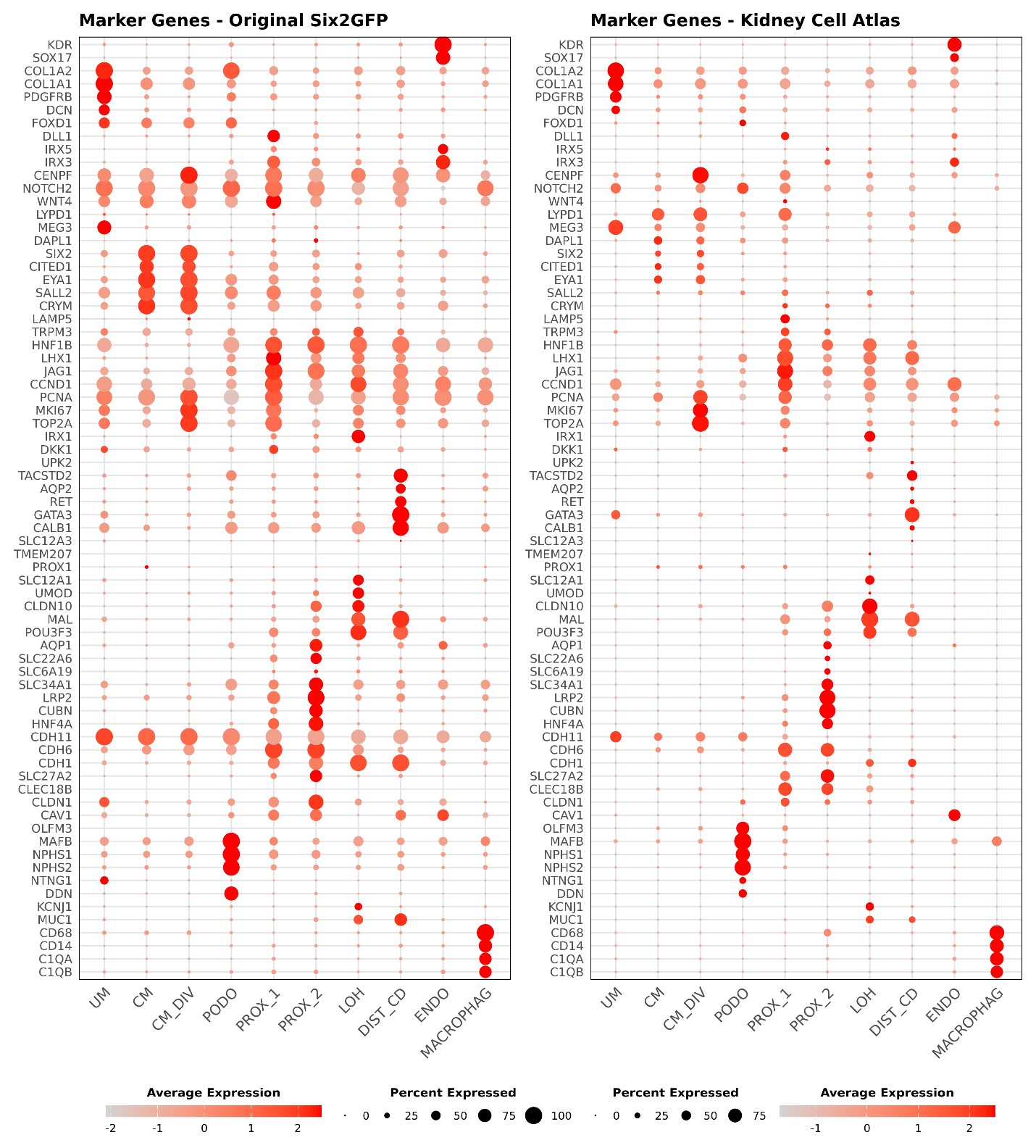

### Figure S3: At the resolution of the 10 major cell types used in our study, cell-label transfer from the mouse fetal kidney to the human fetal kidney yields cell subpopulations whose marker gene expression resembles that of their reference counterparts.

Dot plots showing marker gene expression in the mouse fetal kidney (Six2-GFP transgenic mouse, Wineberg et al. [3], left) and the human fetal kidney (Kidney Cell Atlas [4], right). Dot size indicates the percentage of cells expressing each gene, and color indicates average expression level. Genes were selected from the DevKidCC publication [5] with additional published markers of fetal kidney cell subpopulations [3]. Note that the mouse dataset was generated with the Smart-seq2 protocol (higher coverage, lower dropout) and the human dataset with the 10x protocol (lower coverage, higher dropout). Marker gene expression profiles of corresponding subpopulations resemble each other despite the cross-species and technical differences between the two datasets.

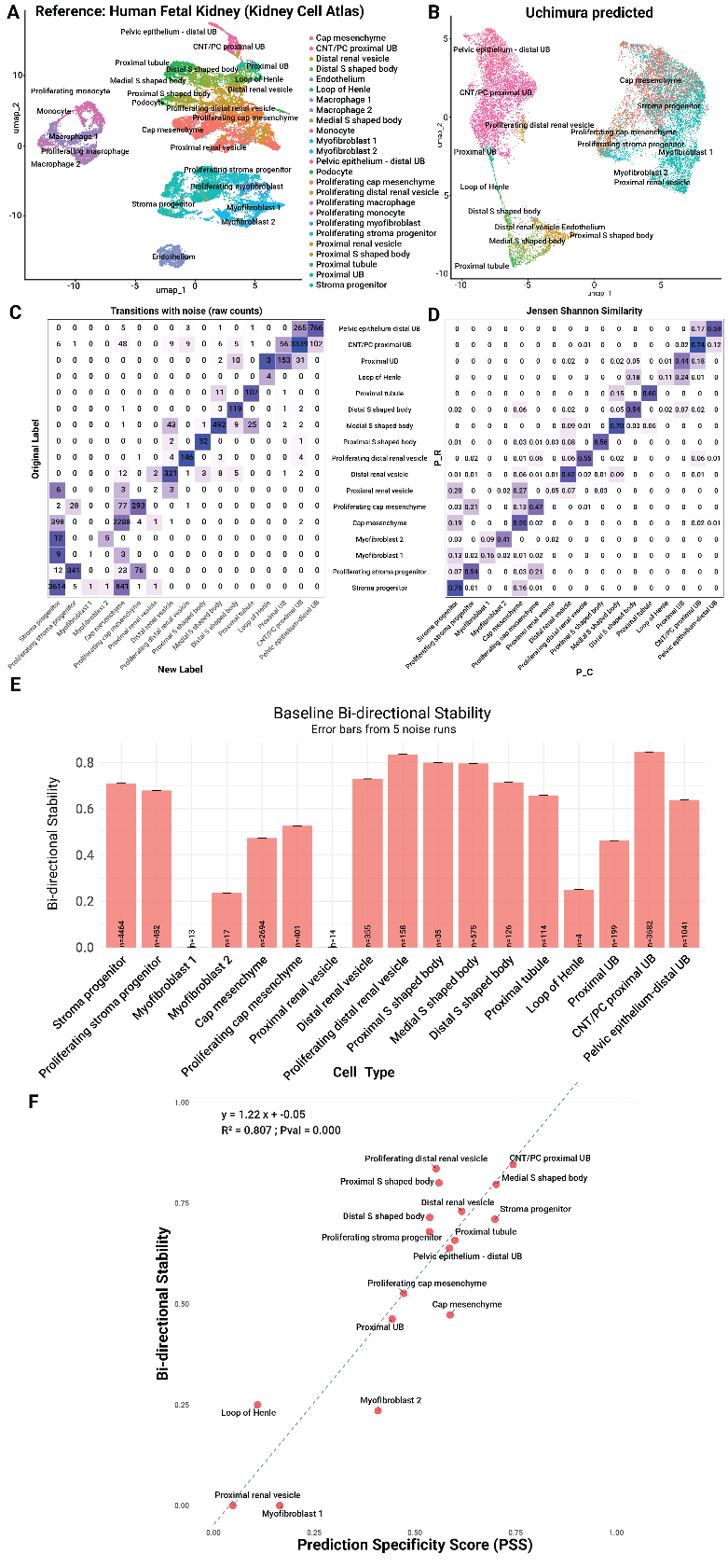

### Figure S4: Cell-label transfer from the human fetal kidney (kidney cell atlas) to the Uchimura et al. iPSC-derived kidney organoid yielded similar results to those obtained when using the mouse fetal kidney as reference.

(A) A UMAP plot of the single-cell RNA-seq reference dataset from the human fetal kidney (kidney cell atlas) [4]. Cell labels were taken from the original publication, in which cells were manually annotated based on known marker genes. (B) A UMAP plot of the single-cell RNA-seq query dataset from Uchimura et al. iPSC-derived kidney organoid [6]. Cells are colored according to their predicted labels. (C) A transition matrix showing the number of cells that switched labels after Poisson noise was applied. Matrix entries are colored as in Figure 2. Only renal cell subpopulations were included. (D) A matrix of Jensen-Shannon similarity scores. Matrix entries are colored as in Figure 2. Only renal cell subpopulations were included. (E) Classification stability scores of renal cell subpopulations, averaged over N=5 noise realizations. (F) A scatter plot of classification stability versus prediction specificity scores for renal cell subpopulations. The two measures show a strong correlation. Similar to observations obtained using the mouse fetal kidney dataset as reference (Figure 3), the Uchimura et al. organoid has high scores for collecting duct lineages (“CNT/PC proximal UB”, i.e., connecting tubule/principal cells derived from the proximal ureteric bud). Note that some cell subpopulations of early epithelial structures (e.g., proximal, medial, and distal S-shaped body) have high scores, likely due to the finer resolution of cell-type annotations in the reference human fetal kidney dataset. Podocytes and proliferating myofibroblasts are not shown since no cells in the query dataset were assigned these labels.
