## Supplementary figures and images for "Robustness measures for cell type classification in single-cell RNA-seq datasets of kidney organoids"

### Figure S1

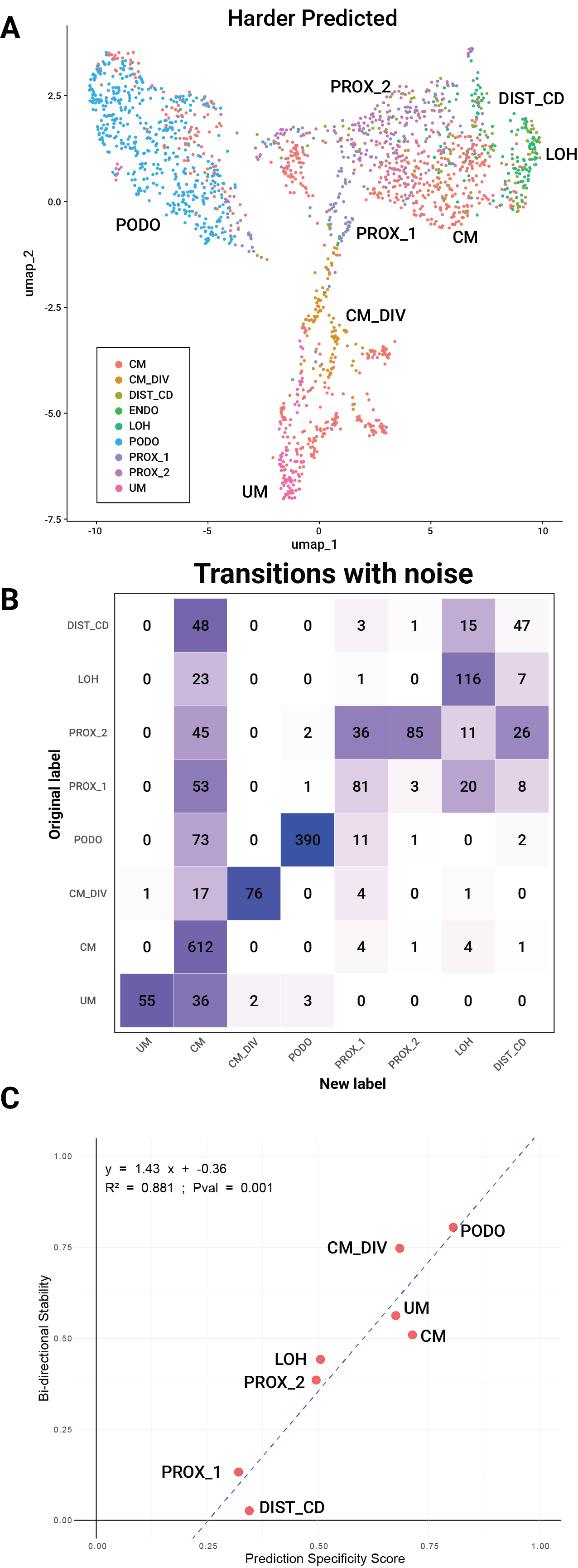

### Figure S2

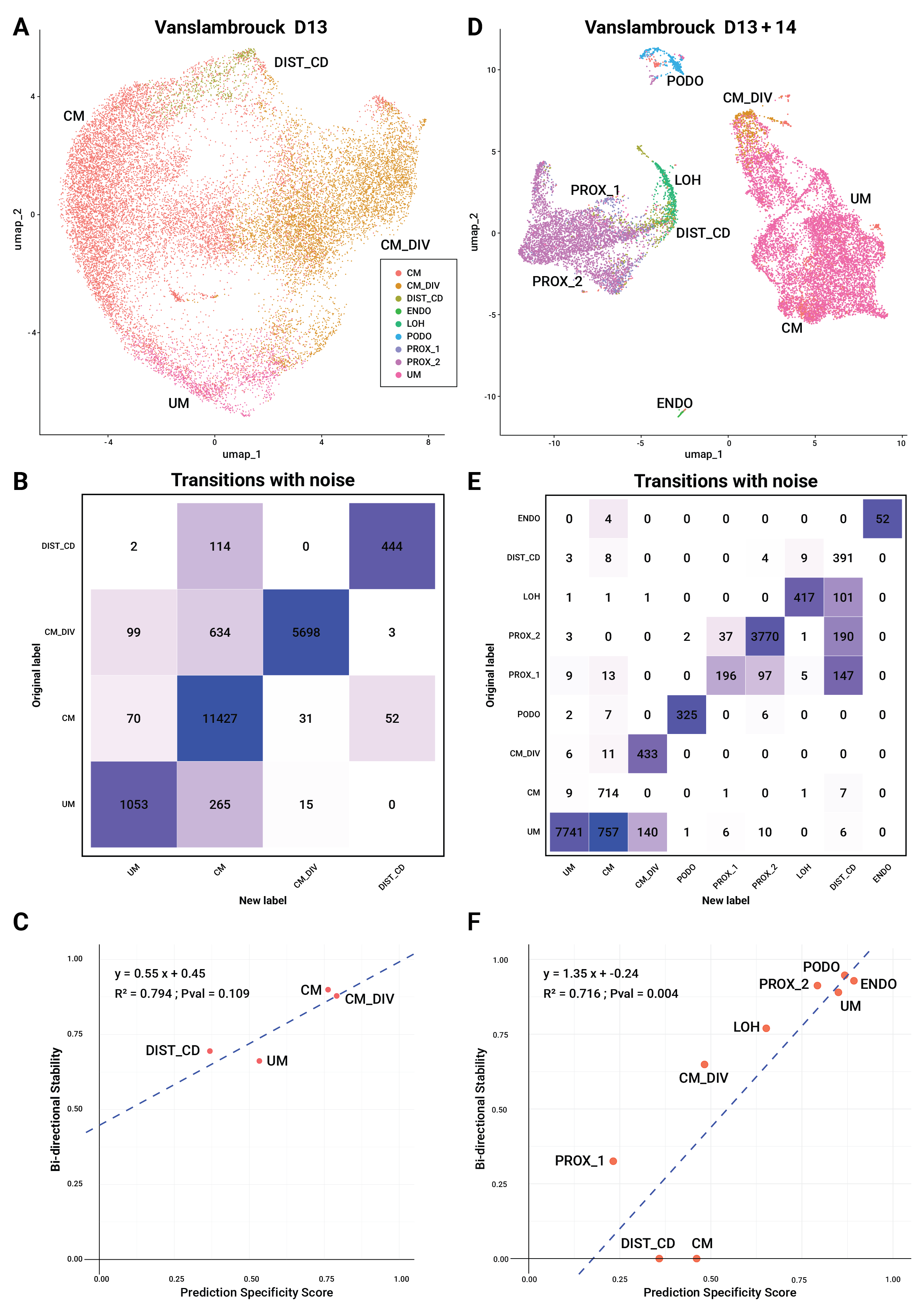

### Figure S3

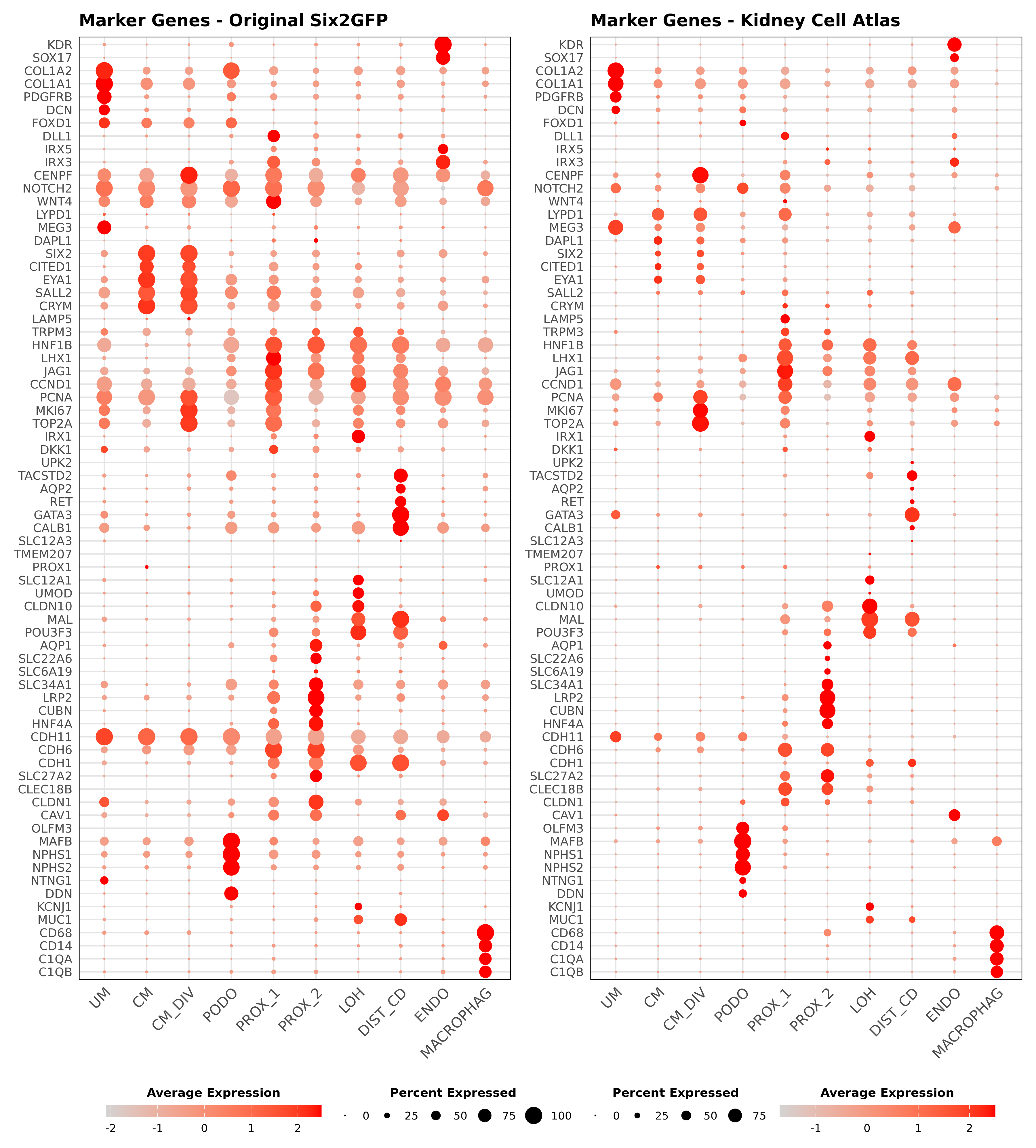

### Figure S4

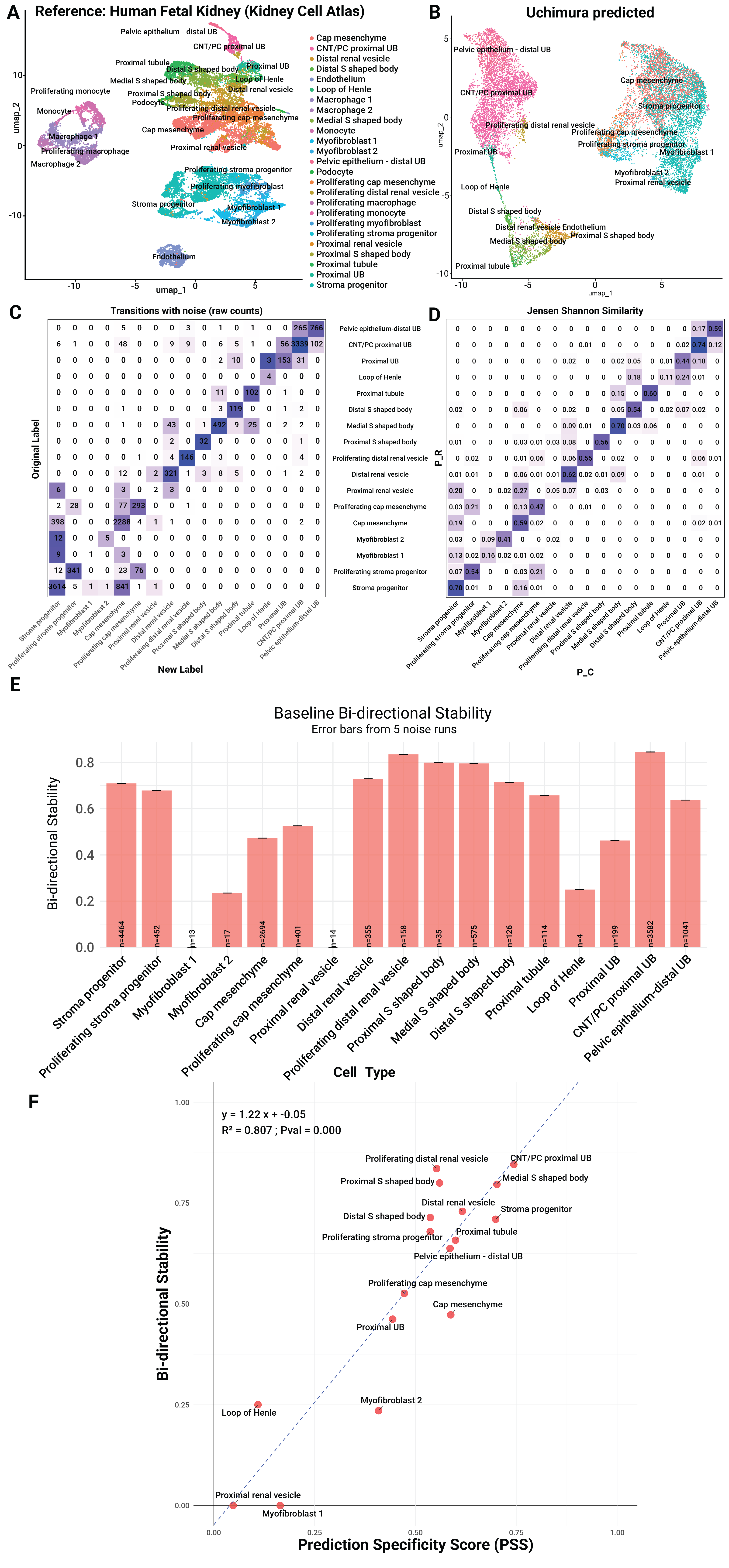
